# A test statistic to quantify treelikeness in phylogenetics

**DOI:** 10.1101/2021.02.16.431544

**Authors:** Caitlin Cherryh, Bui Quang Minh, Rob Lanfear

## Abstract

Most phylogenetic analyses assume that the evolutionary history of an alignment (either that of a single locus, or of multiple concatenated loci) can be described by a single bifurcating tree, the so-called the treelikeness assumption. Treelikeness can be violated by biological events such as recombination, introgression, or incomplete lineage sorting, and by systematic errors in phylogenetic analyses. The incorrect assumption of treelikeness may then mislead phylogenetic inferences. To quantify and test for treelikeness in alignments, we develop a test statistic which we call the tree proportion. This statistic quantifies the proportion of the edge weights in a phylogenetic network that are represented in a bifurcating phylogenetic tree of the same alignment. We extend this statistic to a statistical test of treelikeness using a parametric bootstrap. We use extensive simulations to compare tree proportion to a range of related approaches. We show that tree proportion successfully identifies non-treelikeness in a wide range of simulation scenarios, and discuss its strengths and weaknesses compared to other approaches. The power of the tree-proportion test to reject non-treelike alignments can be lower than some other approaches, but these approaches tend to be limited in their scope and/or the ease with which they can be interpreted. Our recommendation is to test treelikeness of sequence alignments with both tree proportion and mosaic methods such as 3Seq. The scripts necessary to replicate this study are available at https://github.com/caitlinch/treelikeness

## Introduction

A phylogenetic tree is a representation of the relationships between species or individuals. Many estimates of phylogenetic trees implicitly assume that sites in a sequence alignment share the same evolutionary history and conform to a single bifurcating tree. This assumption is called treelikeness. This concept was first introduced by Dress (1984) and was first used to assess how well data fit a tree by Eigen *et al*. (1988). Perfectly treelike alignments are likely to be rare not only due to noise such as sequencing or alignment error but also because biological processes like incomplete lineage sorting (ILS), recombination, or introgression mean even very short alignments may have an evolutionary history that cannot be represented by a single bifurcating phylogeny (Mallet *et al*. 2016; Mendes *et al*. 2019; Scornavacca and Galtier 2017). Although the treelikeness assumption is almost universally made in phylogenetic analyses, it remains rare to test the validity of this assumption. If treelikeness is incorrectly assumed, phylogenetic inferences may be misled (Brown and Thomson 2018), so it is important to test whether the treelikeness assumption holds prior to estimating a phylogenetic tree. Doing so may assist phylogeneticisits in choosing the best combination of data and inference method with which to infer their phylogeny.

Most estimates of phylogenetic trees assume that the data are treelike at some level. Concatenation methods (also known as supermatrix methods) assume that all loci in an alignment share a single evolutionary history. This approach has been criticised as the histories of individual loci may vary dramatically, potentially resulting in incorrect phylogenetic inferences (Shi and Yang 2018; Weisrock *et al*. 2012; Wielstra *et al*. 2014; Wu *et al*. 2018; Zhao *et al*. 2016). Concatenating alignments from loci with different evolutionary histories clearly violates the treelikeness assumption. Coalescent methods improve on the concatenation method by explicitly incorporating non-treelikeness due to ILS. To do so, coalescent methods allow each locus to have a separate tree topology and use the distribution of these topologies to then infer the species tree under a model of ILS. However, these methods still assume that the alignment used to produce each single-locus tree is treelike. Recent work shows that different exons in the same gene often have different evolutionary histories, meaning that in many cases the alignments used to estimate single-locus trees may not be treelike (Mendes *et al*. 2019; Scornavacca and Galtier 2017). As a result, both concatenation and coalescent tree estimation methods may be vulnerable to errors introduced by violation of the treelikeness assumption.

Previous studies have proposed a range of approaches to measuring certain aspects of non-treelikeness. Goldman (1993) developed the first general test for model adequacy in evolutionary models, which simultaneously assess all assumptions of the evolutionary model, including the treelikeness assumption. Unfortunately, as all parameters of the model are tested simultaneously, the treelikeness of the alignment cannot be extracted from the results of this test. Additionally, several methods for visualizing treelikeness have been suggested. Likelihood mapping (Strimmer and von Haeseler 1997) is primarily used to visualise and estimate the phylogenetic signal of an alignment (Baric *et al*. 2003; Salzburger *et al*. 2002; Steiner and Dreyer 2003) but has also been used to assess whether an alignment had a treelike structure (Nadan *et al*. 2003). The *δ* plot method (Holland *et al*. 2002) allows assessment of the treelikeness of an alignment using a mathematical approach based on assessing the treelikeness of all possible quartets of taxa in the alignment. The mean *δ*_q_ value from the *δ* plot method has been used to draw inferences about the overall treelikeness of an alignment, although the interpretation of this value varies: values of 0.11 (Kozak *et al*. 2015) and 0.28 (Short *et al*. 2014) have been suggested to indicate significant non-bifurcating signal, and a value of 0.18 was suggested to indicate that an alignment was treelike (Grimm and Renner 2013).

Phylogenetic split networks have also been used to visualise the treelikeness of an alignment. A split network generalises a phylogenetic tree by representing incompatible phylogenetic signals present in the sequence alignment as additional edges (Huson *et al*. 2010). Compared to a tree, a phylogenetic split network includes more information about the relationships between taxa as it includes conflicting phylogenetic signals and alternate histories (Bryant and Moulton 2004). The treelikeness of alignments can be visually determined based on the number, size and position of parallelograms within the network (Bryant and Moulton 2004; Kennedy *et al*. 2005; Kück *et al*. 2010). Phylogenetic networks provide a very useful visual tool for assessing treelikeness, but they can be somewhat difficult to interpret and there currently exists no framework for comparing the treelikeness of different alignments using networks. Ideally, a test statistic should quantify treelikeness in a way that is comparable across alignments and allow biologists to make informed decisions about which data and methods to use for inferring evolutionary histories.

Here, we introduce the tree proportion for quantifying the treelikeness of a multiple sequence alignment. The tree proportion of an alignment describes the proportion of a phylogenetic network of that alignment that can be represented by a single bifurcating tree. Specifically, it is the proportion of non-trivial split weights of an inferred network that are contained in a bifurcating phylogenetic tree of the same alignment. A split is trivial if one side of the bipartition contains only one taxon, i.e. terminal branches on a phylogeny represent trivial splits, because they are contained in all trees and networks and thus provide no information about treelikeness. Tree proportion ranges from 0 to 1, where a score of 0 indicates that none of the non-trivial splits in the network are represented in the tree. A score of 1 indicates that all of the non-trivial splits in the network are represented in the tree (i.e. the alignment is perfectly treelike). More generally, the better that bifurcating phylogenetic tree represents an alignment, the closer the tree proportion will be to 1.

In addition to providing an intuitive measure of treelikeness, we describe how the tree proportion can be used to ask whether the assumption of treelikeness can be rejected for any given alignment. To do this, we use a parametric bootstrap to simulate treelike datasets with model parameters estimated from the original alignment, and then ask whether the observed tree proportion is surprisingly small relative to the tree proportions observed from the (truly treelike) simulated datasets. This allows us to generate a p-value for the test statistic, where a p-value < 0.05 indicates that the assumption of treelikeness can be rejected for the alignment in question. Finally, using introgression as a framework for simulating alignments of varying treelikeness, we demonstrate how the tree proportion can be used to quantify and test for treelikeness, and we compare its performance to previously suggested methods for estimating treelikeness, as well as certain measures that have been suggested specifically for testing introgression.

## New approaches

### Tree proportion

Tree proportion is defined as follows. For a split network denoted by (S, *λ*) where *S* is the set of non-trivial splits and *λ* is a split weight function, and a phylogenetic tree (T) the tree proportion is calculated as:

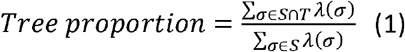

In other words, tree proportion is the proportion of the total weight of non-trivial splits in the network that are represented by the tree. Figure 1 illustrates how to calculate the tree proportion for a simple five-taxon split network and tree.

**Figure 1.**
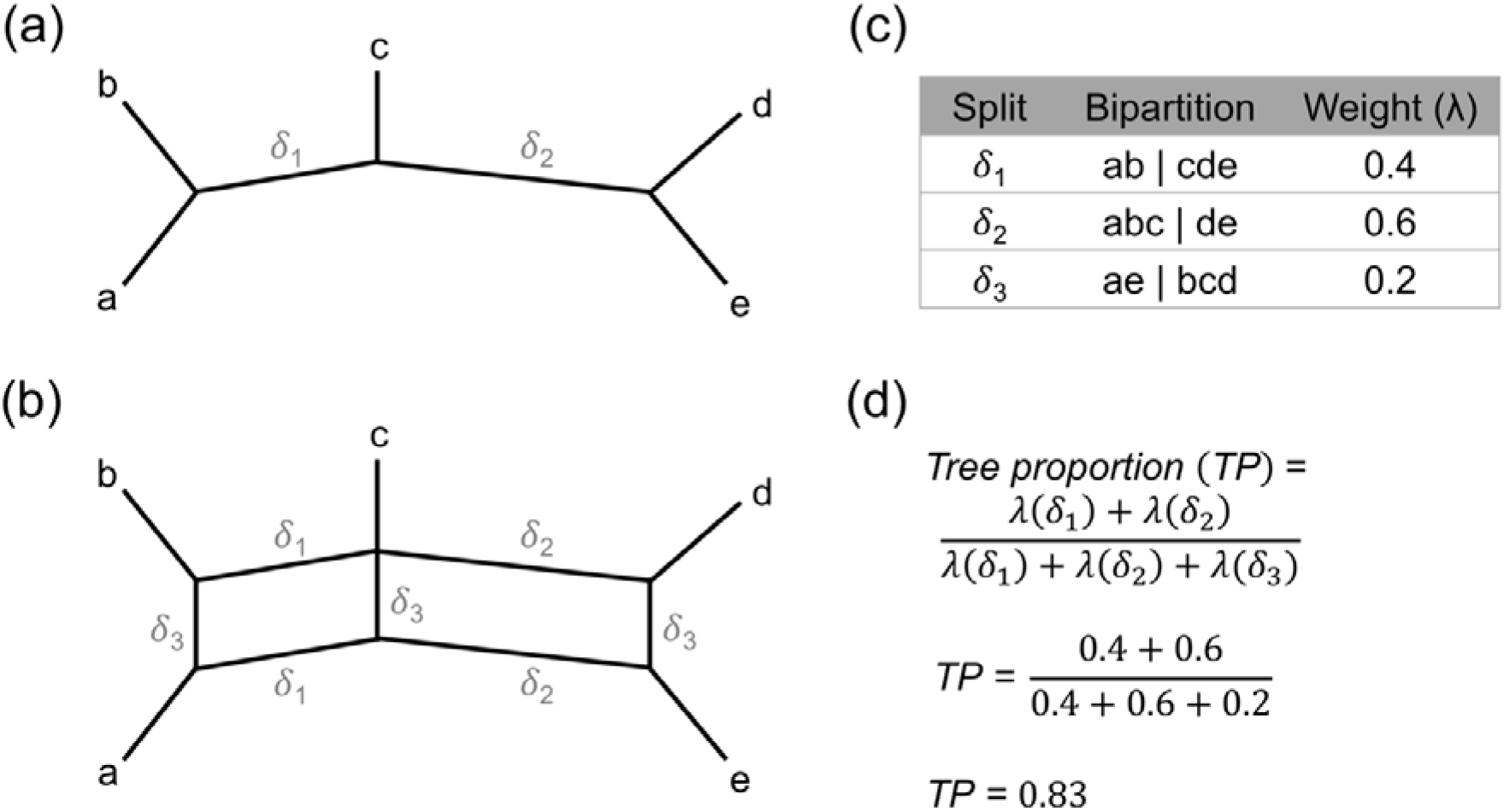
**a.** a simple phylogenetic tree for five taxa. Non-trivial splits are labelled. **b.** A split phylogenetic network for the same five taxa. Non-trivial splits are labelled. **c.** Table showing the bipartition of taxa and weight for each non-trivial split. **d.** Sample calculation of the tree proportion for the alignment with tree **a** and network **b**.

Calculating tree proportion requires both a bifurcating phylogenetic tree and a phylogenetic split network estimated from the same alignment. In principle the split network and the bifurcating phylogenetic tree could be inferred with any method. Indeed, the maximum tree proportion for any given split network can be calculated simply from using the maximum spanning tree of the network as the bifurcating tree. In this study, we used Maximum Likelihood to estimate bifurcating trees for two reasons: (1) Maximum Likelihood is one of the most commonly used methods for tree inference, and (2) Maximum Likelihood naturally allows us to extend our approach to include a parametric bootstrap test because it co-estimates the bifurcating tree and the parameters of a model of molecular evolution. We used NeighborNet (Bryant and Moulton 2004) to estimate the split network. NeighborNet is a distance based agglomerative method for generating replicable and statistically consistent split networks. NeighborNet measures conflict rather than evolutionary history, so the resulting network represents conflicting signals within the alignment.

To calculate the tree proportion of an alignment, we first estimated a NeighborNet network in SplitsTree v4.14.6 (Huson and Bryant 2006). Next, we estimated a maximum likelihood tree for the same alignment using IQ-Tree v2.0 with ModelFinder (Kalyaanamoorthy *et al*. 2017; Minh *et al*. 2020). Finally, we calculated tree proportion in R using code available from https://github.com/caitlinch/treelikeness.

### Parametric bootstrap

Because we do not assume any prior distribution of tree proportion, we rely on a parametric bootstrap procedure to determine whether the tree proportion is significantly lower than would be expected for truly treelike alignments as follows. For a given alignment *D*, we reconstruct a maximum likelihood tree *T_ML_* with the best-fit substitution model *M*. From *T_ML_* and *M* we simulate *n* (n=199 by default) alignments *D*_1_,…, *D_n_*. From each *D_i_* we reconstruct a maximum likelihood tree *T_i_* and a NeighborNet network *S_i_* and use *T_i_* and *S_i_* to calculate the tree proportion *TP_i_*. We calculate the statistics TP_1_,…,*TP_n_*. The p-value is then computed as the fraction of *TP_i_* greater than or equal to TP of the original alignment.

## Results

### Decreased treelikeness due to increasing proportions of introgressed DNA

Of the six test statistics for treelikeness we compared, tree proportion (R^2^ = 0.863) and mean *δ*_q_ (R^2^ = 0.818) showed the strongest correlations with the proportion of introgressed DNA (Figure 2). Both tree proportion and mean *δ*_q_ were strongly correlated with the proportion of introgressed DNA whether the simulated introgression was reciprocal or non-reciprocal (Supplementary Figure 1: Tree proportion R^2^ = 0.863 and 0.704; mean *δ*_q_ R^2^ = 0.818 and 0.902). Tree proportion was strongly correlated with the proportion of introgressed DNA regardless of the simulated tree depth (Supplementary Figure 2, all R^2^ > 0.692). Mean *δ*_q_ showed strong correlations on tree depths up to 0.5 (Supplementary Figure 2, all R^2^ > 0.705) but a much weaker correlation when the simulated tree depth was 1.0 (Supplementary Figure 2, R^2^ = 0.344). The strength of the correlations between the proportion of introgressed DNA and the other four test statistics was highly variable and never higher than 0.511 under any simulation conditions (Figure 2, Supplementary Figures 1 and 2).

**Figure 2:**
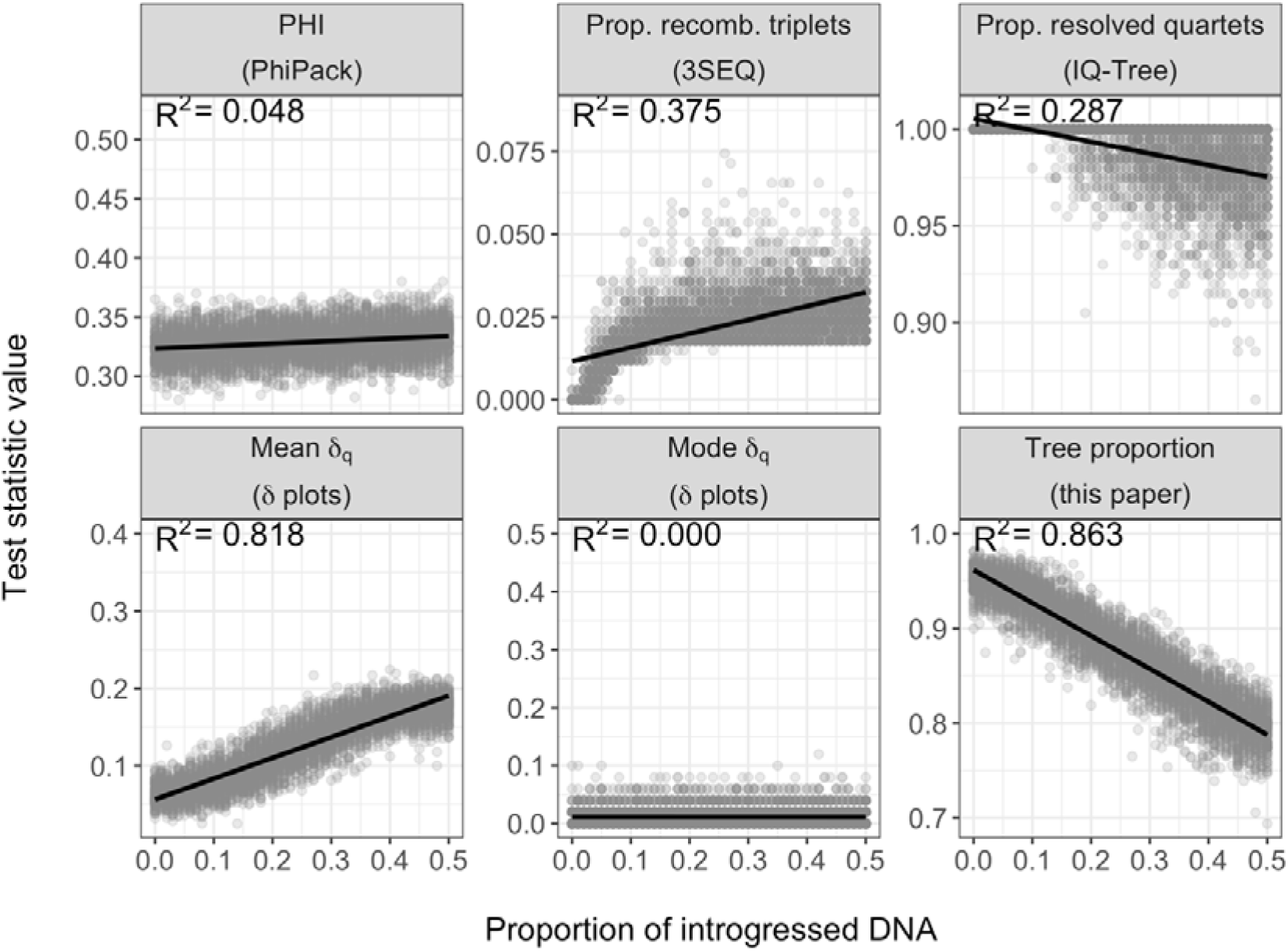
Test statistic values for increasing proportions of introgressed DNA from 0 to 0.5 in 0.01 increments with one close, non-reciprocal introgression events and tree depth of 0.5 (n = 100). Each point represents the test statistic value for a single simulated alignment. Prop. recomb. triplets is the proportion of recombinant triples. Prop. resolved quartets is the proportion of resolved quartets.

The ability of each test to statistically reject treelikeness under simulated introgression events varied greatly (Figure 3). PHI and 3SEQ had the highest power to reject non-treelike alignments, and the tests successfully detect 100% of introgression events after the proportion of introgressed DNA reached 0.2 and 0.1 respectively (Figure 3). Mean *δ*_q_ and tree proportion have intermediate results. At a proportion of introgressed DNA of 0.5, these tests detect 98% and 99% respectively of the alignments containing introgression (Figure 3). Proportion of resolved quartets and mode *δ*_q_ failed as statistical tests and did not successfully detect introgression events for any tree depth or event type (Figure 3, Supplementary Figures 3 and 4). Test results were very similar regardless of whether introgression was simulated as a reciprocal or a non-reciprocal event (Supplementary Figure 3). All test statistics except proportion of resolved quartets and mode δ_q_ have acceptable false positive rates (i.e., a significant result in approximately 5% of tests when there is no introgression in the simulation, shown as 0 on the x-axis of each panel in Figure 3). Both 3SEQ and PHI have less power to detect introgression at lower tree depths. For a tree depth of 0.05 substitutions per site and at a proportion of introgressed DNA of 0.5, PHI and 3SEQ correctly identify 96% and 95% of introgression events respectively (Supplementary Figure 4). Conversely, tree proportion has less power to detect introgression at higher tree depths. For a tree depth of 1 substitution per site at a proportion of introgressed DNA of 0.5, tree proportion correctly identifies just over a third (35%) of introgression events (Supplementary Figure 4).

**Figure 3:**
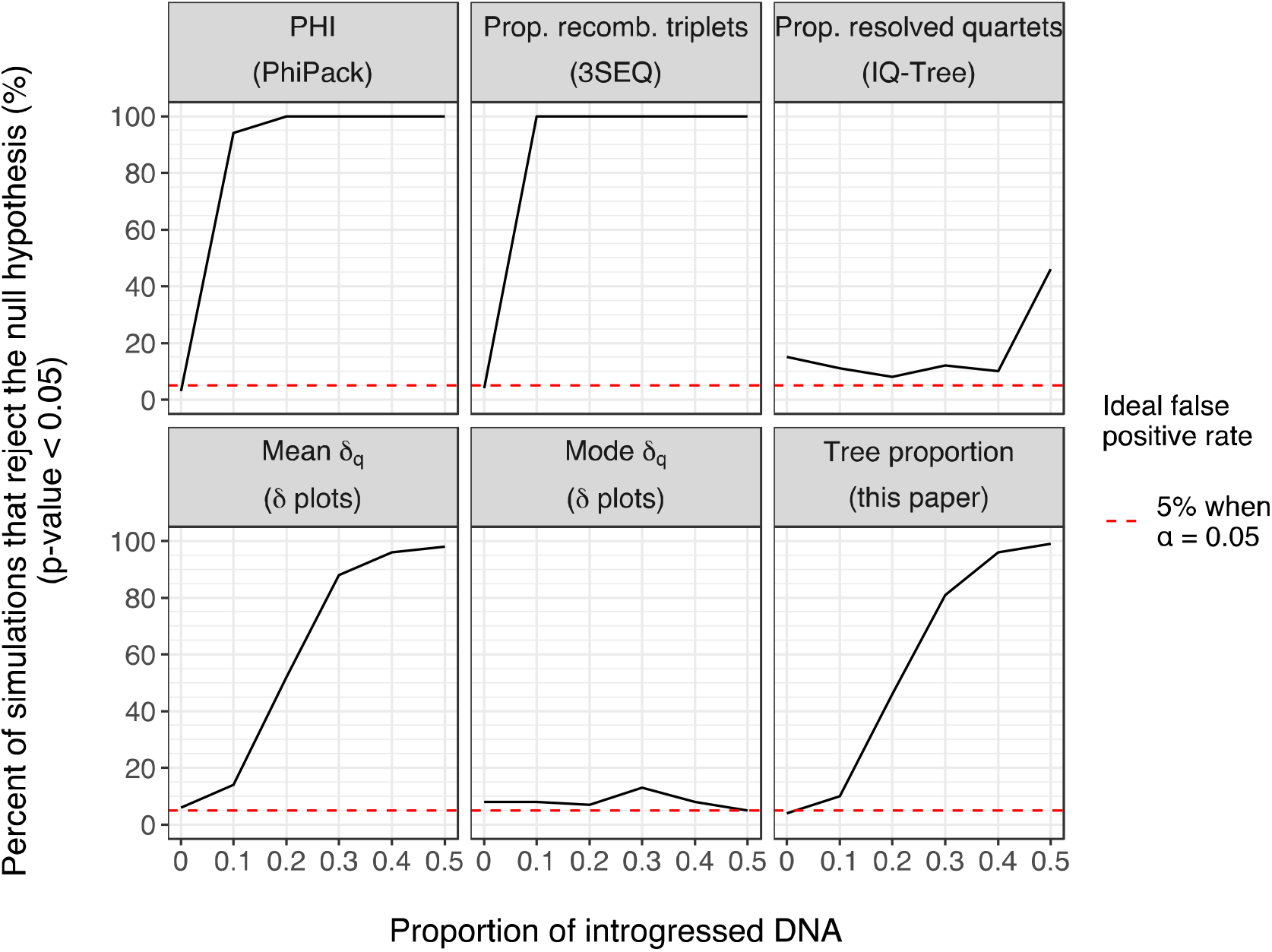
Percentage of simulated alignments that reject the null hypothesis as the proportion of introgressed DNA increases for all six test statistics. Simulated alignments had a single close, non-reciprocal introgression event and tree depth was 0.5. Each line represents the number of alignments (out of 100 replicates) that reject the null hypothesis of treelikeness. Prop. recomb. triplets is the proportion of recombinant triples. Prop. resolved quartets is the proportion of resolved quartets.

### Decreased treelikeness due to increasing number of introgression events

Of the six test statistics we compared, only 3SEQ and tree proportion revealed clear decreases in treelikeness as the number of introgression events increased (Figure 4). Encouragingly, tree proportion responded similarly for all tree depths and whether or not the simulated recombination was reciprocal or non-reciprocal (Supplementary Figures 5 and 6). 3SEQ test statistic values were similar across reciprocal and non-reciprocal recombination events (Supplementary Figure 5), but the range of values is dependent on tree depth (Supplementary Figure 6). The PHI test statistic only identified nonreciprocal introgression events (Supplementary Figure 5), and was more strongly correlated to the number of introgression events at higher tree depths (Supplementary Figure 6). The proportion of resolved quartets, mean *δ*_q_ and mode *δ*_q_ test statistic values showed at best weak correlations with the number of introgression events, regardless of the simulation conditions (Figure 4; Supplementary Figures 5 and 6).

**Figure 4:**
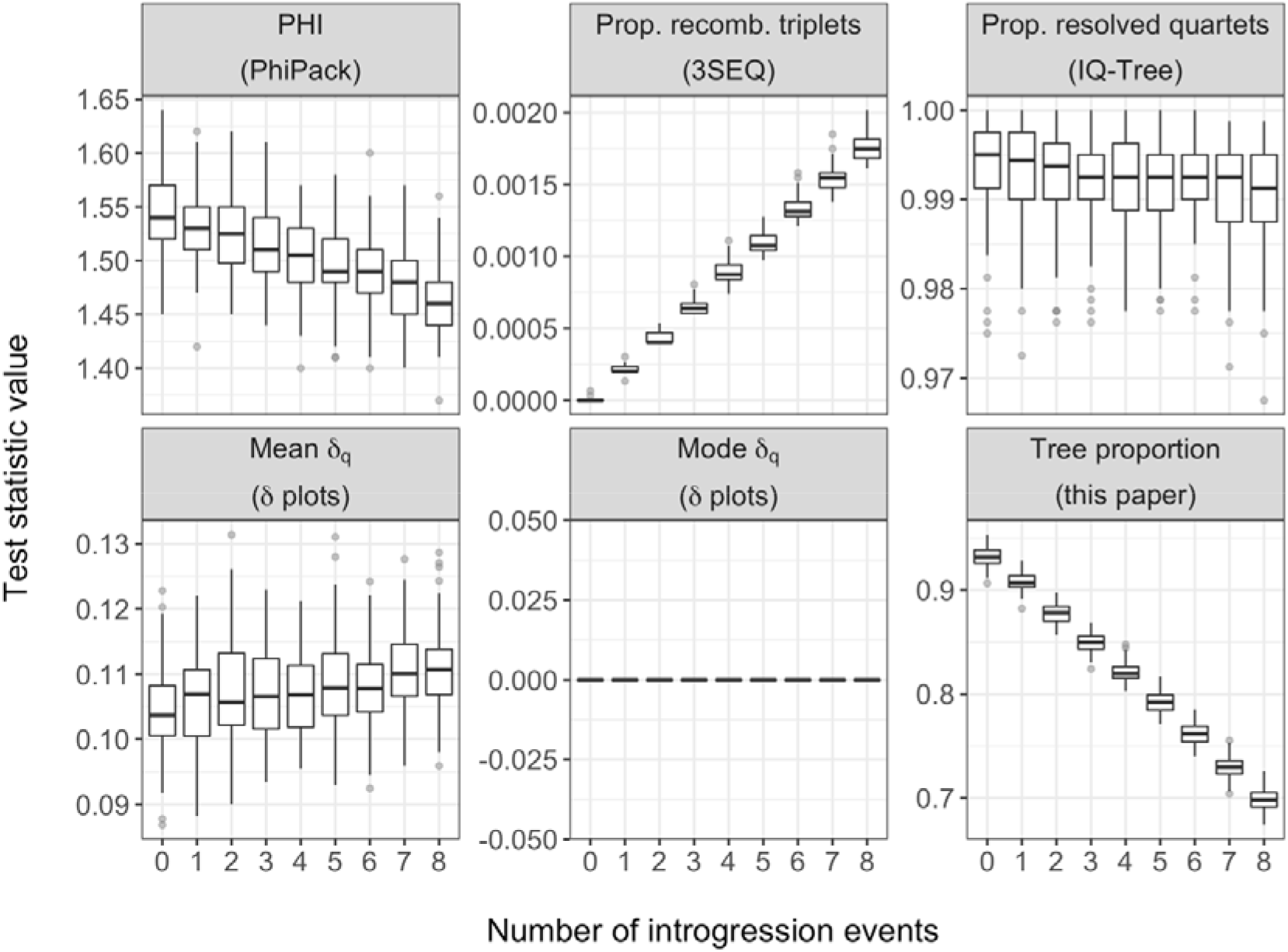
Test statistic values for number of non-reciprocal introgression events and tree depth of 0.5 (n = 100). Each box shows the distribution of test statistic values at a certain number of introgression events. The whiskers extend to the closest observed value no more than 1.5 times the interquartile range away from the box. Points represent outliers. Prop. recomb. triplets is the proportion of recombinant triples. Prop. resolved quartets is the proportion of resolved quartets.

Three test statistics performed well as statistical tests to reject treelikeness in the presence of multiple introgression events (Figure 5). 3SEQ, tree proportion and PHI had the highest power to reject non-treelike alignments, as the tests successfully detected 100% of alignments containing introgression after 1, 2, and 3 events respectively (Figure 5). These results were similar for nonreciprocal and reciprocal events (Supplementary Figure 7). The best performing test statistic was tree proportion, which behaved similarly for all event types and tree depths (Supplementary Figures 7 and 8). 3SEQ performed well at tree depth of 0.5 substitutions per site (Figure 5) but showed very poor performance at the lowest tree depth of 0.05 substitutions per site (Supplementary Figure 8). PHI also performed well at tree depth of 0.5 substitutions per site (Figure 5), but its performance dropped below that of tree proportion at low tree depths (Supplementary Figure 8). The other three statistics (proportion of resolved quartets, mean *δ*_q_ and mode *δ*_q_) were unable to reject treelikeness for the simulated introgressed alignments at any tree depth or event type (Figure 5, Supplementary Figures 7 and 8). Only PHI and tree proportion had acceptable false positive rates (i.e., a significant result in approximately 5% of tests when there is no introgression in the simulation, shown as 0 on the x-axis of each panel in Figure 5).

**Figure 5:**
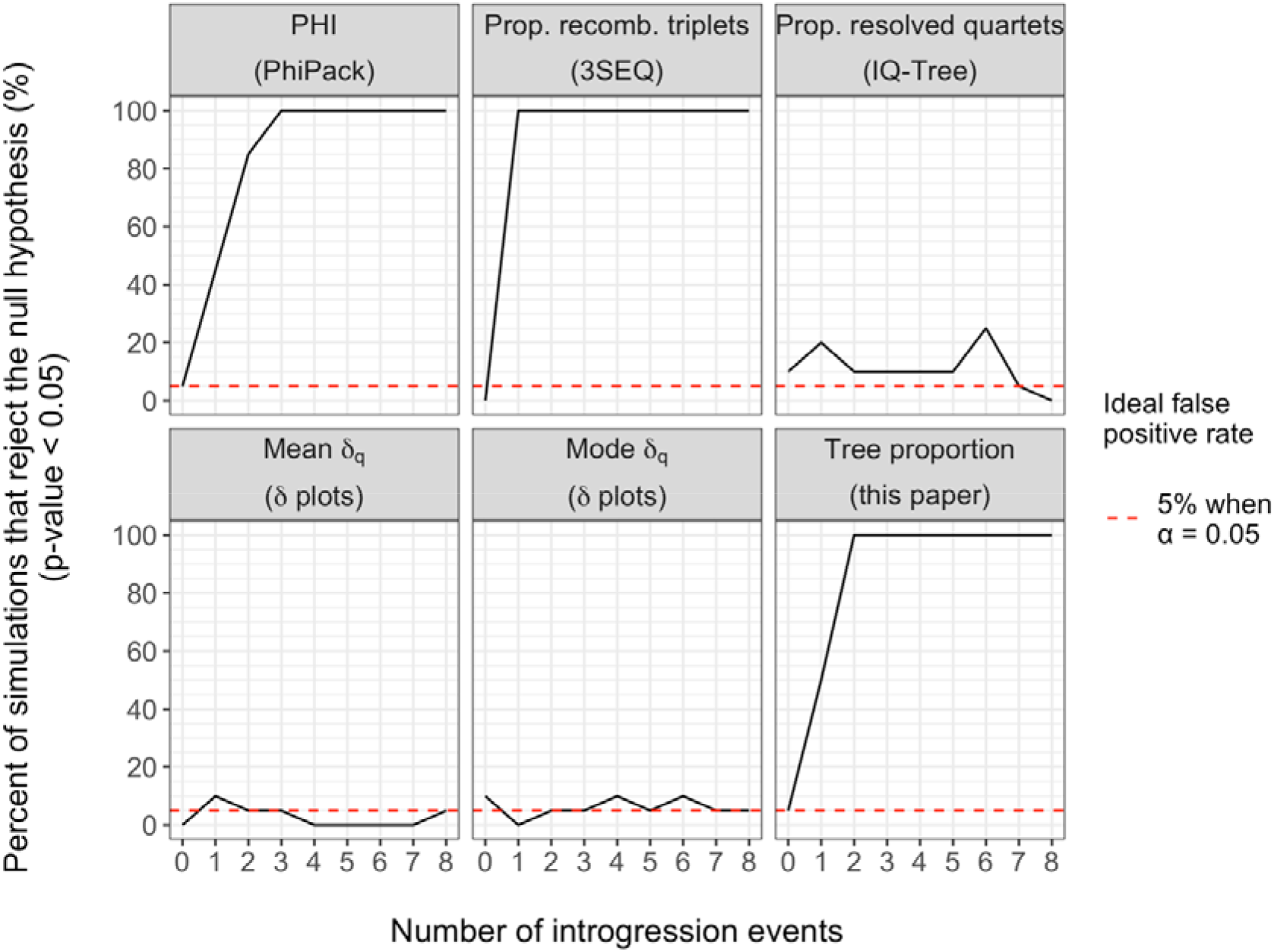
Percentage of simulated alignments that reject the null hypothesis as the number of non-reciprocal introgression events increases for all six test statistics. Simulated alignments had a tree depth of 0.5. Each line represents the number of alignments (out of 100 replicates) that reject the null hypothesis of treelikeness. The red dotted line represents the ideal false positive rate of 5% when *α* = 0.05. Prop. recomb. triplets is the proportion of recombinant triples. Prop. resolved quartets is the proportion of resolved quartets.

## Discussion

In this study, we introduce the tree proportion as a way of measuring treelikeness, and testing (with a parametric bootstrap) whether a single bifurcating phylogenetic tree is sufficient to explain the evolutionary history of an alignment. Importantly for a proposed measure of treelikeness, tree proportion values are easy to interpret: a value of 1 corresponds to a perfectly treelike alignment (i.e. one whose evolutionary history can be perfectly explained by a single bifurcating phylogenetic tree), and as treelikeness reduces the tree proportion will decrease towards zero. These properties mean that the tree proportion can be used to directly compare the treelikeness of different alignments.

We use a suite of simulations to compare the tree proportion to five other tests, both as a measure of treelikeness and as a statistical test that can be used to reject treelikeness for a given alignment. Our results show that the tree proportion is a very useful measure of treelikeness: under a huge range of simulation conditions tree proportion consistently declines in concert with declines in the treelikeness of the alignment. When used as a statistical test to ask whether an alignment can reject treelikeness, the six test statistics we compare have varied success at detecting multiple causes of decreased treelikeness, and no one test performed the best across all simulation conditions. Tree proportion, 3SEQ (Lam *et al*. 2018) and PHI (Bruen *et al*. 2006) performed well as statistical tests rejecting treelikeness under simulated introgression events, and we found that at least one of these tests detected almost every simulated recombinant alignment in the majority of cases. It is perhaps unsurprising that PHI and 3SEQ performed well: both tests are designed to detect recombinant sequences of exactly the type we simulated. However, introgression is just one example of a biological event that reduces the treelikeness of an alignment, and our simulations show that both PHI and 3SEQ performed poorly in certain simulation conditions. As a result, we suggest that statistical tests of treelikeness for empirical alignments would be best served by combining a phylogenetic approach such as tree proportion with a mosaic-based test such as 3SEQ. The tree proportion test works in a wide variety of conditions, and produces an easy-to-interpret test statistic. However, in many conditions (such as if the non-treelikeness is caused by introgression and the tree depth is above a certain threshold) 3SEQ has much more power to detect non-treelikeness. Using both tests therefore provides the most generality and power across all possible causes of non-treelikeness that may impact phylogenetic analyses.

Tree proportion joins a growing group of tests for absolute model adequacy. Penny *et al*. (1992) wrote that a fundamental criterion for a scientific method is that the data must be able to reject the model, a requirement that is rarely met in phylogenetics (Brown and Thomson 2018). Most phylogenetic analyses proceed with only relative tests of model adequacy (such as ModelFinder (Kalyaanamoorthy *et al*. 2017) which selects the best model from a pre-defined set of a models) or no test (Cui *et al*. 2013; de Souza *et al*. 2018; Grismer *et al*. 2018; Grybchuk *et al*. 2018; Kang *et al*. 2014; Lei and Dong 2016; Pearce *et al*. 2017; Tay *et al*. 2017). As model violation is widespread across phylogenetic datasets (Naser-Khdour *et al*. 2019), phylogenetic analyses may benefit if absolute tests for model adequacy are performed prior to tree estimation. Tree proportion builds on the absolute test for model accuracy developed by Goldman (1993). While Goldman’s test encompasses all assumptions of the tree and model which are used to calculate the likelihood in phylogenetic analyses, the tree proportion assesses a subset of model assumptions, and asks specifically to what extent a single bifurcating tree is adequate for explaining the evolutionary history of a given alignment.

The PHI test has been widely used to detect recombination in phylogenetic alignments (Cabanne *et al*. 2008; Croll and Sanders 2009; Croucher *et al*. 2015; D’Horta *et al*. 2011; Faria *et al*. 2016; Harris *et al*. 2012; Joly and Bruneau 2006; Ogura *et al*. 2009; Pinho *et al*. 2008; Tian *et al*. 2012; Weinert *et al*. 2009). The PHI test performed well in our simulations and detected almost all introgression events under all simulation conditions. Similarly, Bruen *et al*. (2006) and Haubold *et al*. (2013) found the PHI test accurately detected recombination in simulated coalescent data and in empirical datasets including bacteria, fungi and virus DNA, and animal mtDNA. However, the PHI test performs poorly when sequence diversity is low (less than 10%), when alignments are short and/or when the number of taxa is low (less than 10) (Bruen *et al*. 2006; White *et al*. 2013; White and Gemmell 2009). In previous simulations where the power of the PHI test was low, the sequence diversity ranged from 0.01 to 1.25 x 10^-3^ (Bruen *et al*. 2006; White *et al*. 2013; White and Gemmell 2009). The diversity in our simulated alignments was much higher than this, and therefore our simulations contained sufficient informative sites and incompatibilities for the test to perform well. The PHI test is a powerful and accurate test for recombination when the sequence diversity and number of taxa are sufficiently large, and is conservative and results in false negatives when they are not.

Likelihood mapping and δ plots allow visualisation of the phylogenetic content of a sequence alignment, but we show here that they do not perform well as test statistics for treelikeness. The mean δ value has been used to quantify the treelikeness of an alignment, with values from 0 to 0.2 generally interpreted as highly treelike (Coiro and Barone Lumaga 2018; Dashper *et al*. 2017; Grimm and Renner 2013; Kozak *et al*. 2015; Meier-Kolthoff and Göker 2019; Short *et al*. 2014; Stanborough *et al*. 2018). However, the mean δ responds inconsistently to causes of decreased treelikeness, so a low mean δ value does not necessarily indicate that an alignment is treelike (e.g., the mean δ value was 0.048 for one of our simulations in which the alignment contained 8 introgression events). Similarly, the proportion of resolved quartets from likelihood mapping has been interpreted as an indicator of the treelikeness of an alignment, with values from 0.8 and up interpreted as treelike (Buesa *et al*. 2002; Elena *et al*. 2001; Li *et al*. 2017; Morgan *et al*. 2014; Nadan *et al*. 2003; Pitra *et al*. 2002; Salemi *et al*. 2000; Shi *et al*. 2012; Verbruggen and Theriot 2008). Likelihood mapping displays the phylogenetic content of an alignment by plotting the treelikeness of individual quartets. However, our results show that this correlates very poorly with the overall treelikeness of an alignment (e.g., the proportion of resolved quartets was 1.0 for some of our simulations in which the alignment contained 8 introgression events). The proportion of resolved quartets for our simulations is high – the minimum value was 70% (Supplementary Figure 2), but the majority of simulations had a proportion of resolved quartets above 85%, despite most of them containing significant non-treelikeness such as multiple introgression events involving a large fraction of the alignment. The reason that the proportion of resolved quartets remains high in these simulations is that each introgression event involves only a few taxa, meaning only a small proportion of quartets are affected and the contribution to the proportion of resolved quartets is low. As a result of these limitations, we recommend using these methods for visual assessment of phylogenetic information, but not for quantifying or testing for treelikeness.

While the parametric bootstrap approach we propose here is designed to isolate non-treelikeness from other signals in the data, this separation will not always be perfect. A statistically significant result from the parametric bootstrap indicates a significant difference in test statistic values between the original alignment and the bootstrap replicates. In our simulations, the only difference between the original alignment and the bootstrap replicates was that the former included introgression events. The same will not be true for empirical alignments, as the models we use for empirical data are gross oversimplifications of the true underlying process (e.g. (Song *et al*. 2010) (Lemmon and Moriarty 2004)). As a result, the parametric bootstrap may return a significant result if other types of model violation lead to differences in the treelikeness between the empirical and the bootstrap-replicate alignment. Given that model violation is widespread and common within phylogenetic datasets (Naser-Khdour *et al*. 2019), we suggest that a significant result from the parametric bootstrap should be interpreted as likely, but not certain, to be caused by non-treelikeness in the empirical alignment. Regardless of the cause, a significant result from the parametric bootstrap should be cause for concern, and perhaps warrant further investigation of the offending alignment.

## Materials and Methods

### Simulation approach

Introgression is a potential source of non-treelikeness that is known to mislead phylogenetic inferences (Posada and Crandall 2002; Wiens 1998). However, it is hard to account for with current methods. Introgression provides a framework wherein different types and amounts of non-treelikeness can be simulated on a linear scale (Posada and Crandall 2002). In this study, we use introgression as a framework for simulating non-treelike alignments in order to compare new and existing measures and tests for treelikeness. Here, we compare tree proportion with two tests for introgression: the Pairwise Homoplasy Index (PHI) (Bruen *et al*. 2006) and 3SEQ (Boni *et al*. 2007; Lam *et al*. 2018). We also applied two existing methods that have been used to test for treelikeness in previous studies: 8 plotting (Holland *et al*. 2002) and likelihood mapping (Strimmer and von Haeseler 1997).

PHI measures the minimum number of convergent mutations on any tree to describe the genealogy of a pair of sites. PHI is a widely used test for recombination or introgression and was previously found to outperform other similar tests (Bruen *et al*. 2006). We calculated the PHI value and p-value for the each alignment using PhiPack (Bruen 2005).

The second test, 3SEQ, attempts to calculate the number and location of introgression events for a given alignment by testing each triplet of sequences using a hypergeometric random walk to determine if one sequence is the child of the other two (Boni *et al*. 2007; Lam *et al*. 2018). 3SEQ has been shown to perform well in simulations (Boni *et al*. 2007; Lam *et al*. 2018). We used the 3SEQ implementation (Lam *et al*. 2018) to calculate the number of recombinant triplets, number of recombinant sequences and p-value for each alignment.

We also applied both the *δ* plotting (Holland *et al*. 2002) and likelihood mapping (Strimmer and von Haeseler 1997) methods, which have been previously used to estimate treelikeness of an alignment. To obtain a test statistic for the *δ* plotting method, we applied the delta.plot function in the R package ape v5.4-1 (Paradis *et al*. 2004) to the distance matrix for each alignment and calculated the mean and mode *δ*_q_ value. The mean or mode *δ*_q_ will be between 0 and 1, where larger values are less treelike. We used the likelihood mapping implementation in IQ-Tree (Minh *et al*. 2020) with the number of quartets to sample set to 25 times the number of taxa, and took the test statistic to be the number of fully-resolved quartets. This test statistic is a proportion, with a value of 1 indicating that every quartet sampled was treelike. The test statistic value decreases as the quartets become less treelike. The p-values for both the *δ* plotting and likelihood mapping methods were calculated using a parametric bootstrap.

### Simulating multiple sequence alignments with introgression

To simulate alignments with introgression, we extended the two-tree simulation approach described in Posada and Crandall (2002) and shown in Figure 6. This method uses forward time phylogenetic simulations to simulate alignments which mimic those that would be produced by introgression events and allows for control over the placement and timing and of introgression events. In principle, one alignment can be simulated along a tree and then a portion of DNA from one species replaced by a portion of DNA from a second species. Here, we achieve the same results by simulating DNA along two trees and then concatenating the two sequences to mimic the result of introgression. This approach provides a simple and flexible framework for simulating introgression on multiple sequence alignments (Posada and Crandall 2002).

**Figure 6:**
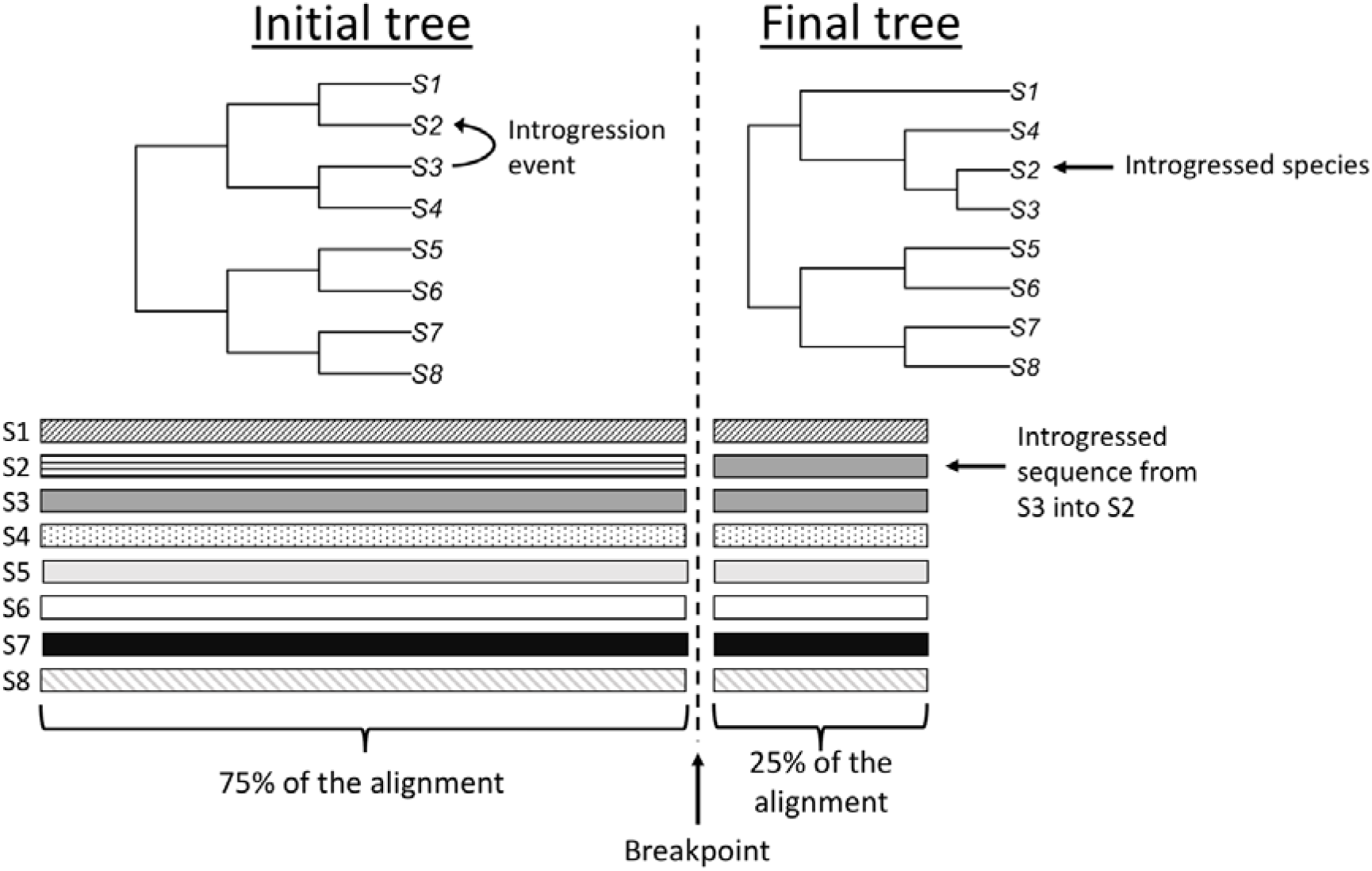
Simulation of a mosaic multiple sequence alignment with an introgression event, adapted from Posada and Crandall (2002). The initial sequence (n% of the final sequence, here n = 75%) is simulated along a balanced 8-taxon tree. The introgression event shown here consists of genetic material from S3 overwriting the original sequence in S2 (as shown by the arrows). In the final tree, the introgression event has occurred, moving the position of S2 in the phylogeny. 25% ((1 - n) %) of the alignment is then simulated along this tree. Two trees are needed to explain the evolutionary history of this alignment, violating the treelikeness assumption.

We used this framework to simulate datasets under two scenarios of varying treelikeness: increasing proportion of introgressed DNA and increasing number of introgression events. Firstly, we investigated the effect of increasing the proportion of introgressed DNA (i.e. proportion of final tree) by simulating sequence alignments on a balanced 8-taxon tree from 0 – 50% introgressed DNA sequence in 1% intervals. 10 replicates were conducted for each set of simulation parameters. We applied the following test statistics to each alignment: tree proportion, *δ* plots, likelihood mapping, PHI test and 3SEQ. Due to the high computational expense of the parametric bootstrap, our tree proportion test was calculated only for the proportion of introgressed DNA from 0% – 50% in 10% intervals (6 intervals total). Secondly, we investigated the effect of increasing the number of introgression events by simulating 0 to 8 introgression events on a 32-taxon balanced tree. A 32-taxon balanced tree has *δ* balanced subtrees, each consisting of two clades with two species each. Each introgression event takes place within one subtree, allowing from 0 to *δ* simultaneous events. For this set of simulations, we fixed the proportion of introgressed DNA at 50%. We performed 100 replicates of each set of simulation parameters. Five test statistics were applied to each alignment as above. The tree proportion test was only performed for the first ten replicates of each set of simulation parameters due to the high computational load.

Other simulation parameters were as follows. All simulations were repeated for reciprocal and non-reciprocal introgression events. A non-reciprocal event, in which DNA is introgressed unidirectionally from one lineage into another, is shown in Figure 6. In a reciprocal event, there is a bidirectional exchange of genetic material between two species. For all simulations, we fixed the sequences length to 1300 base pairs (the average length of a transcript in eukaryotes, from Xu *et al*. (2006)), and the model of substitution to the Jukes-Cantor model (Jukes and Cantor 1969). We simulated four substitution rates (in substitutions per site), 0.05, 0.1, 0.5 and 1, to simulate varying rates of molecular evolution across simulations. The total number of simulated alignments was 4080 for the first set of simulations and 6800 for the second set.

The scripts for this analysis were written in R v3.6.3 (R Core Team 2020) using the packages ape v5.4-1 (Paradis *et al*. 2004), ggplot2 v3.3.2 (Wickham 2016), phangorn v2.5.5 (Schliep 2011), phytools v0.7-20 (Revell 2012), seqinr v3.6.1 (Charif and Lobry 2007), stringr v1.4.0 (Wickham 2019) and TreeSim v2.4 (Stadler 2017). Code to replicate all simulations is available from https://github.com/caitlinch/treelikeness. Results from the simulations containing test statistic and statistical test results for all six tests are available in the article and in its online supplementary material.

## Supporting information

Supplementary Files README

## Acknowledgements

The authors would like to thank Barbara Holland, Lindell Bromham, David Gordon and Rod Peakall for their comments and advice. This work was supported by the Australian Research Council grant no. DP-200103151 to R.L. and B.Q.M.

