## Supplementary Figures for "A test statistic to quantify treelikeness in phylogenetics"

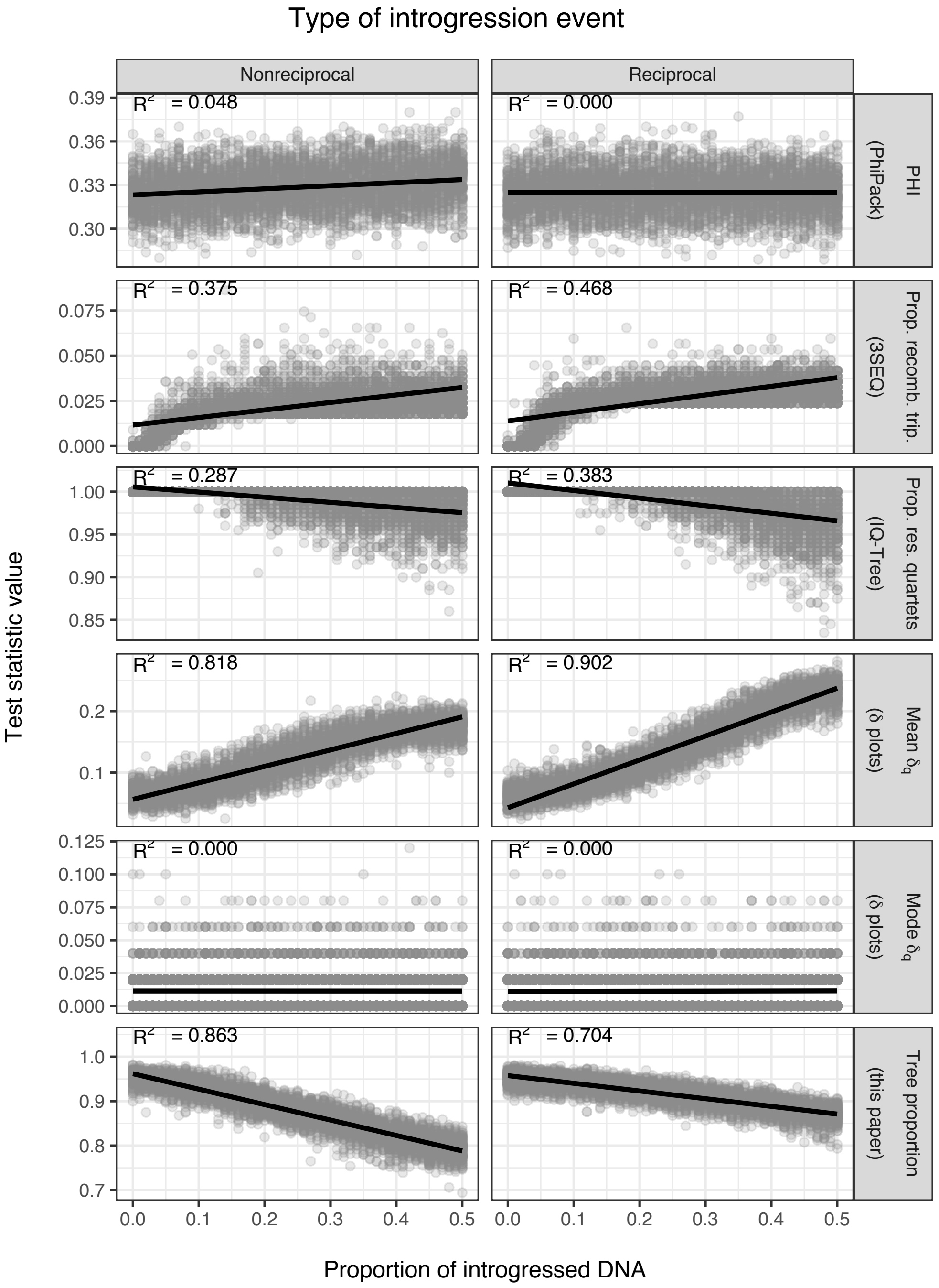


**Supplementary Figure 1:** Changes in test statistic values for a balanced 8-taxon taxon tree with tree depth of 0.5 substitutions per site, with one reciprocal or non-reciprocal introgression event and increasing proportion of introgressed DNA.

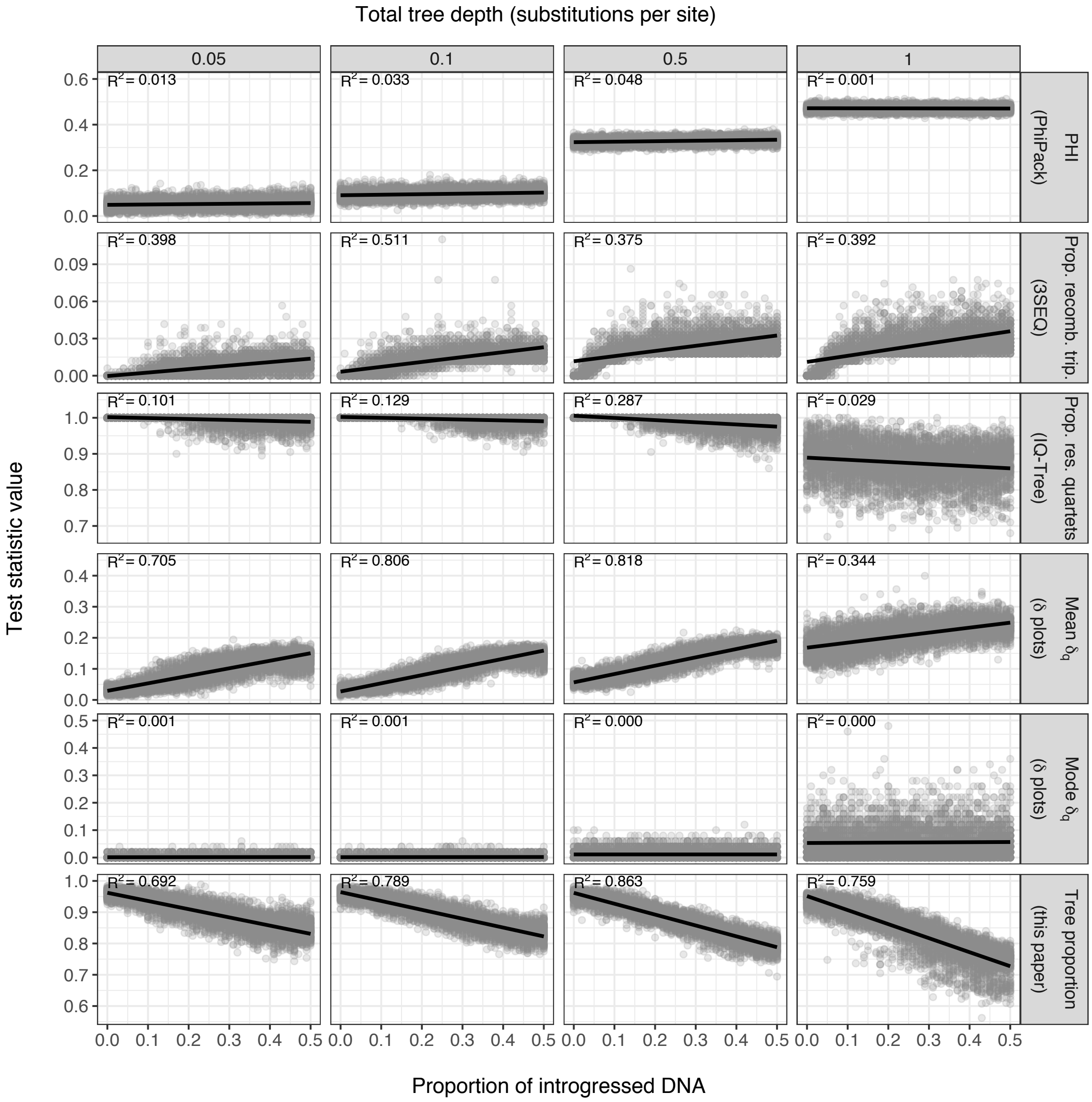


**Supplementary Figure 2:** Changes in test statistic values for a balanced 8-taxon taxon tree with tree depth of 0.5 substitutions per site and one non-reciprocal introgression event, and increasing proportion of introgressed DNA, for four tree depths from 0.05 to 1 (in substitutions per site).

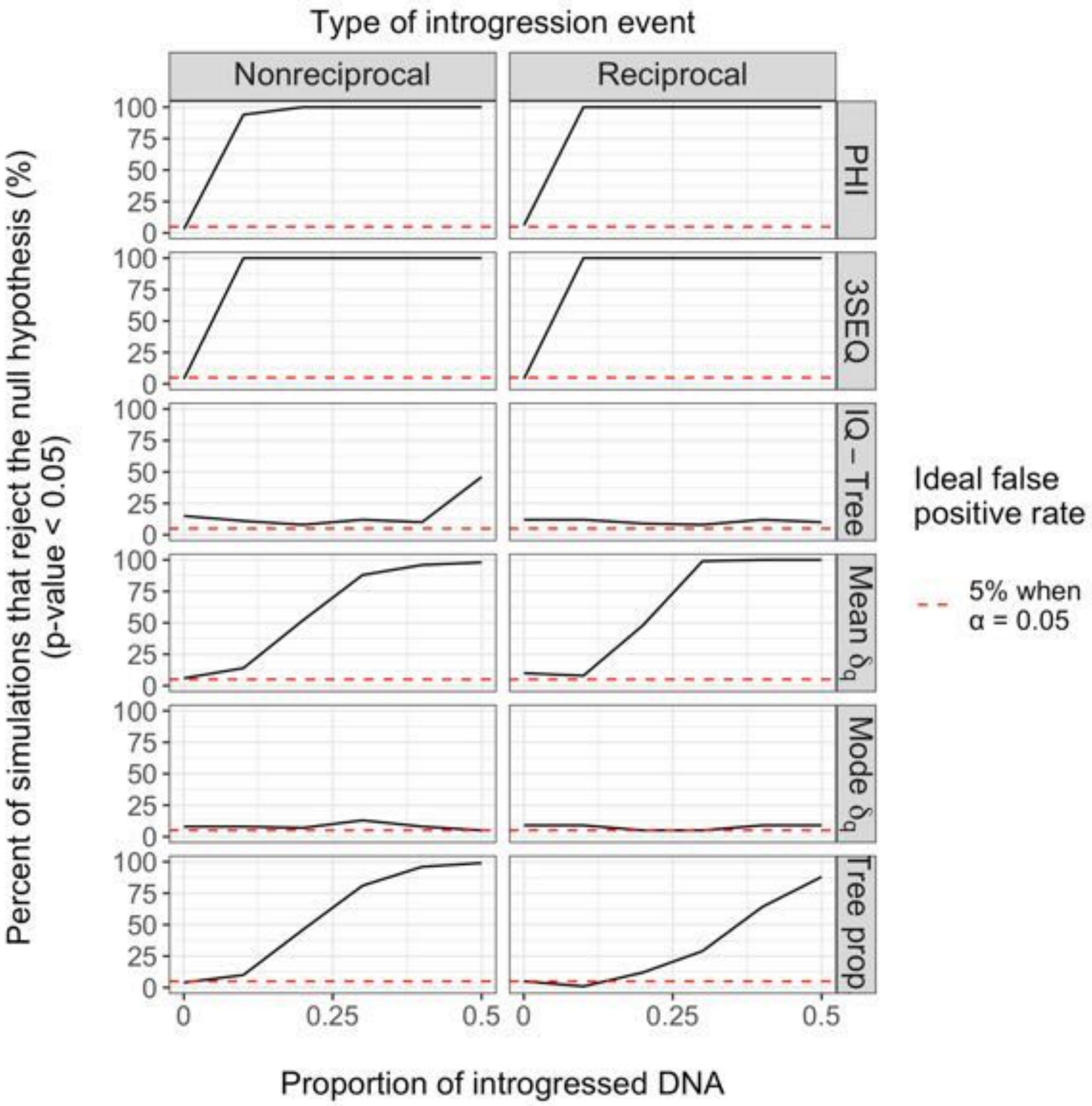


**Supplementary Figure 3:** p-values for 1 nonreciprocal introgression event on a balanced 8 taxon tree with a tree depth of 0.5 substitutions per site as the proportion of introgressed DNA increases from 0 to 50%. Each line represents the number of alignments (out of 100 replicates) that reject the null hypothesis of treelikeness. Results for each event type are shown in columns. Each row shows results from a different test statistic. PHI is the p-value from the PHI test (implemented using PhiPack), 3SEQ is the p-value from the 3SEQ test, IQ-Tree is the p-value calculated using a parametric bootstrap from the proportion of resolved quartets (Likelihood Mapping), Mean $\delta_q$ and Mode $\delta_q$ are the p-values from the $\delta$ plotting method (calculated using a parametric bootstrap), and Tree prop. is the p-value for the tree proportion (calculated using a parametric bootstrap).

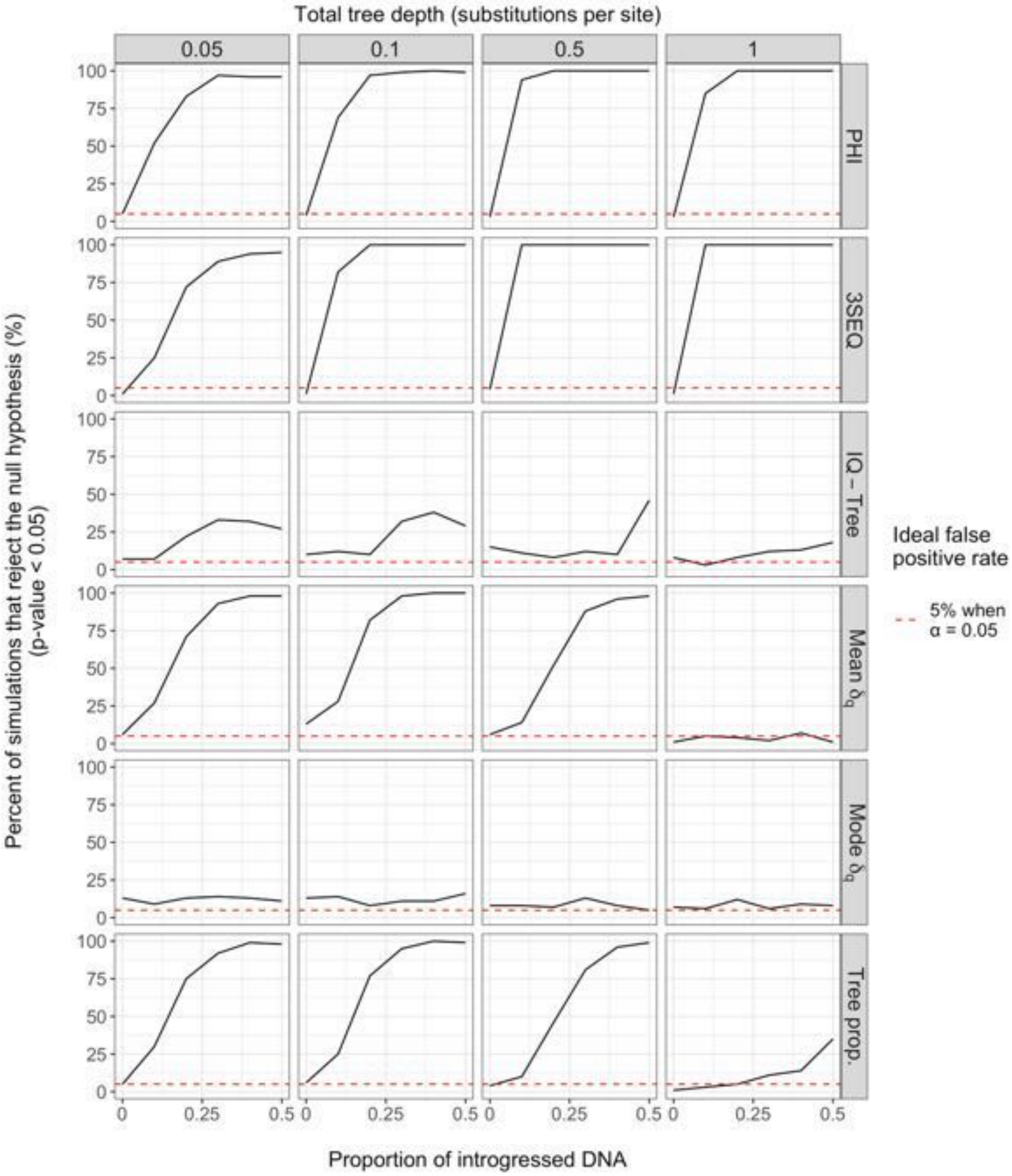


**Supplementary Figure 4:** p-values for 1 nonreciprocal introgression event on a balanced 8 taxon tree with a tree depth of 0.5 substitutions per site as the proportion of introgressed DNA increases from 0 to 50%. Each line represents the number of alignments (out of 100 replicates) that reject the null hypothesis of treelikeness. Each column is a tree depth from 0.05 to 1 in substitutions per site. Each row shows results from a different test statistic. PHI is the p-value from the PHI test (implemented using PhiPack), 3SEQ is the p-value from the 3SEQ test, IQ-Tree is the p-value calculated using a parametric bootstrap from the proportion of resolved quartets (Likelihood Mapping), Mean $\delta_q$ and Mode $\delta_q$ are the p-values from the $\delta$ plotting method (calculated using a parametric bootstrap), and Tree prop. is the p-value for the tree proportion (calculated using a parametric bootstrap).

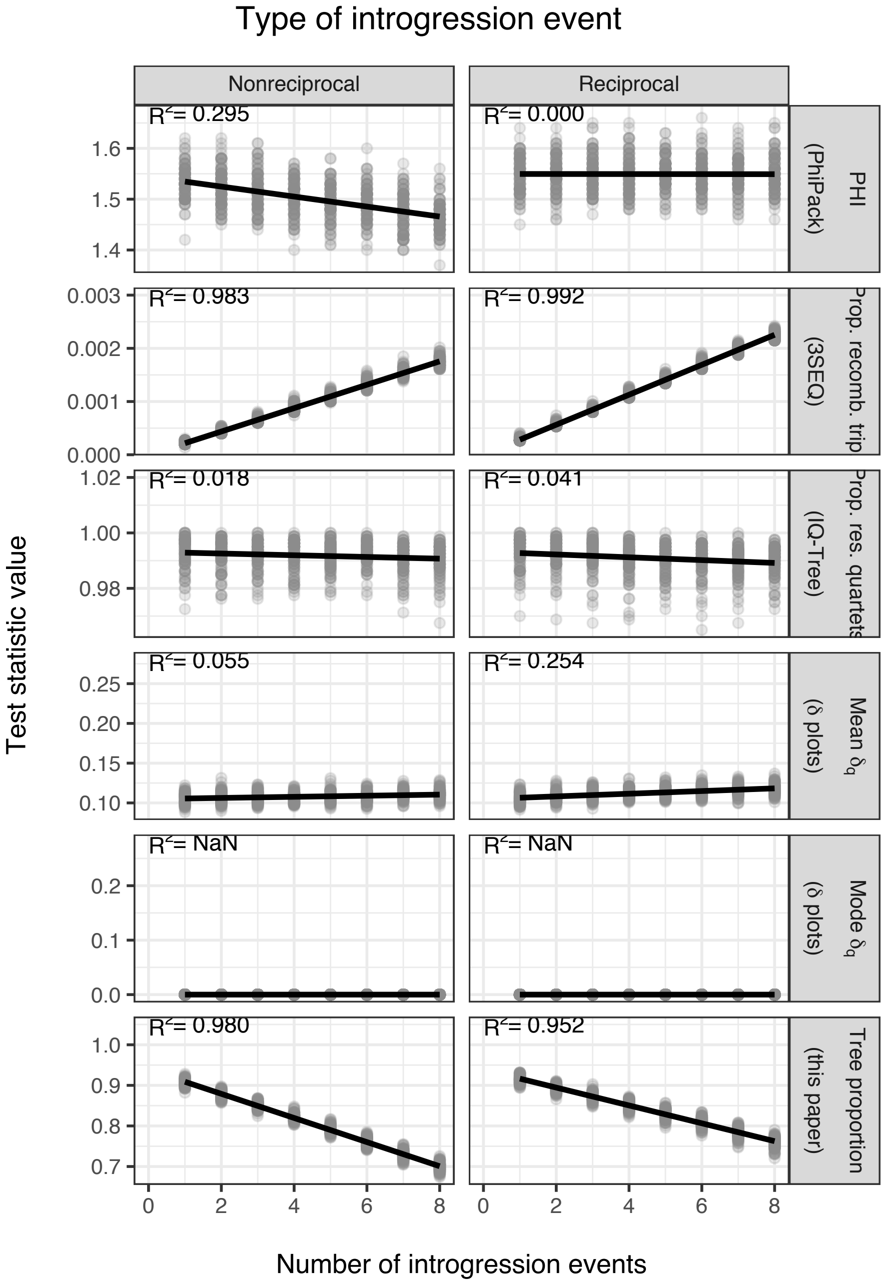


**Supplementary Figure 5:** Changes in test statistic values for a balanced 32-taxon taxon tree with tree depth of 0.5 substitutions per site and 50% introgressed DNA, with increasing number of reciprocal or non-reciprocal introgression events.

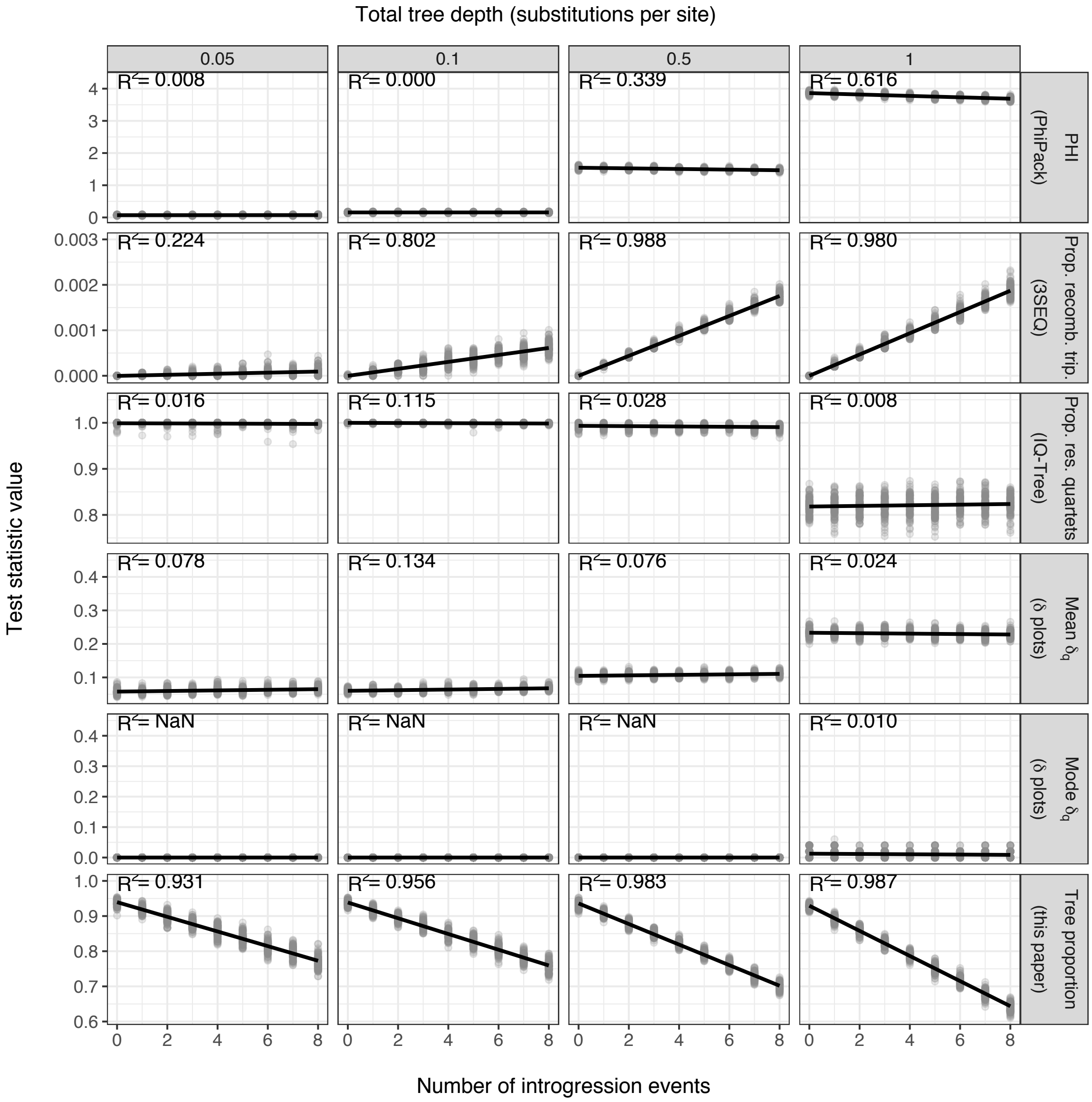


**Supplementary Figure 6:** Changes in test statistic values for a balanced 32-taxon taxon tree with increasing number of non-reciprocal introgression events and 50% introgressed DNA, for four tree depths from 0.05 to 1 (in substitutions per site).

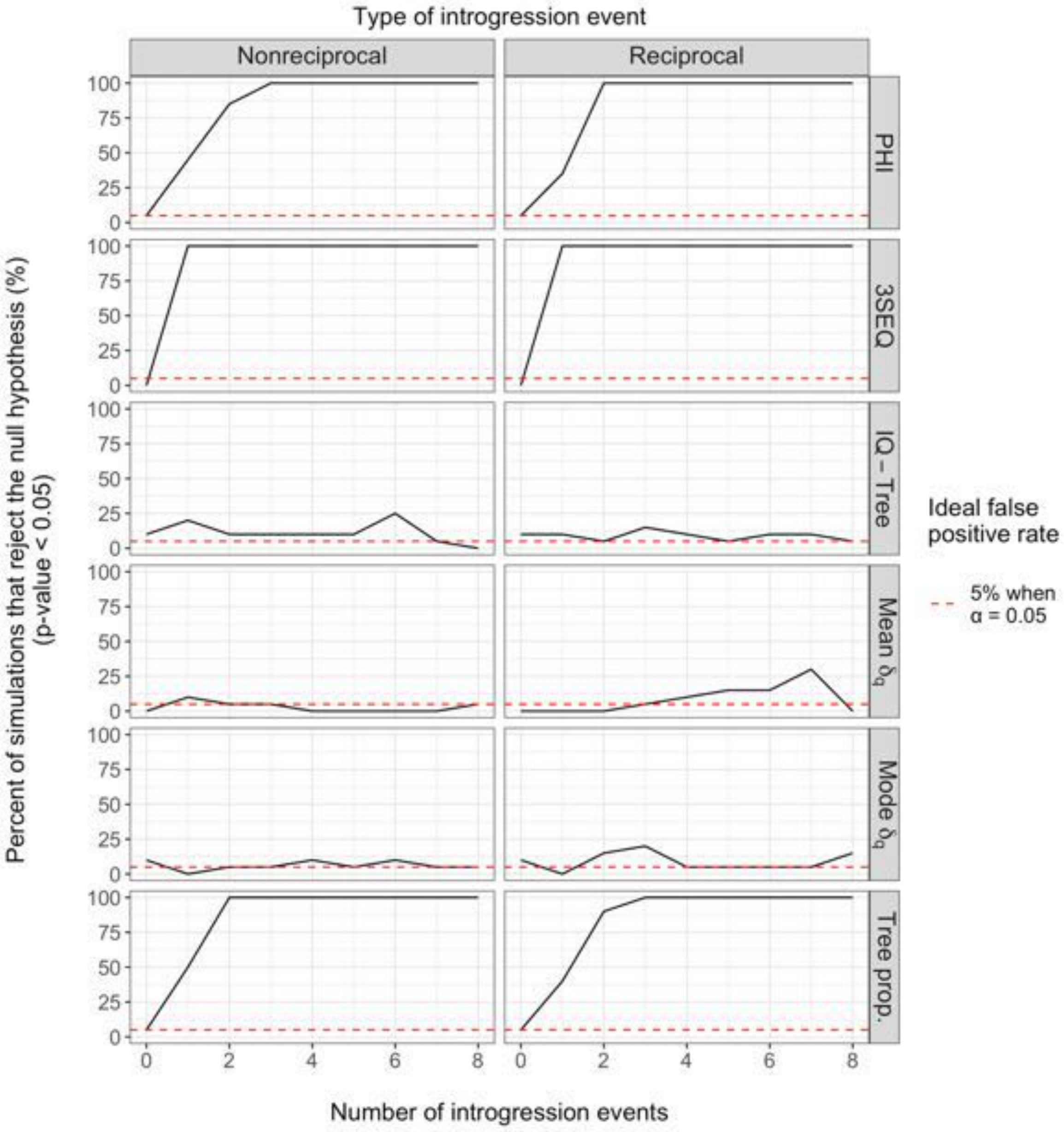


**Supplementary Figure 7:** p-values for an increasing number of nonreciprocal introgression events on a balanced 32-taxon tree with a tree depth of 0.5 substitutions per site. Each line represents the number of alignments (out of 100 replicates) that reject the null hypothesis of treelikeness. Results for each event type are shown in columns. Each row shows results from a different test statistic. PHI is the p-value from the PHI test (implemented using PhiPack), 3SEQ is the p-value from the 3SEQ test, IQ-Tree is the p-value calculated using a parametric bootstrap from the proportion of resolved quartets (Likelihood Mapping), Mean $\delta_q$ and Mode $\delta_q$ are the p-values from the $\delta$ plotting method (calculated using a parametric bootstrap), and Tree prop. is the p-value for the tree proportion (calculated using a parametric bootstrap).

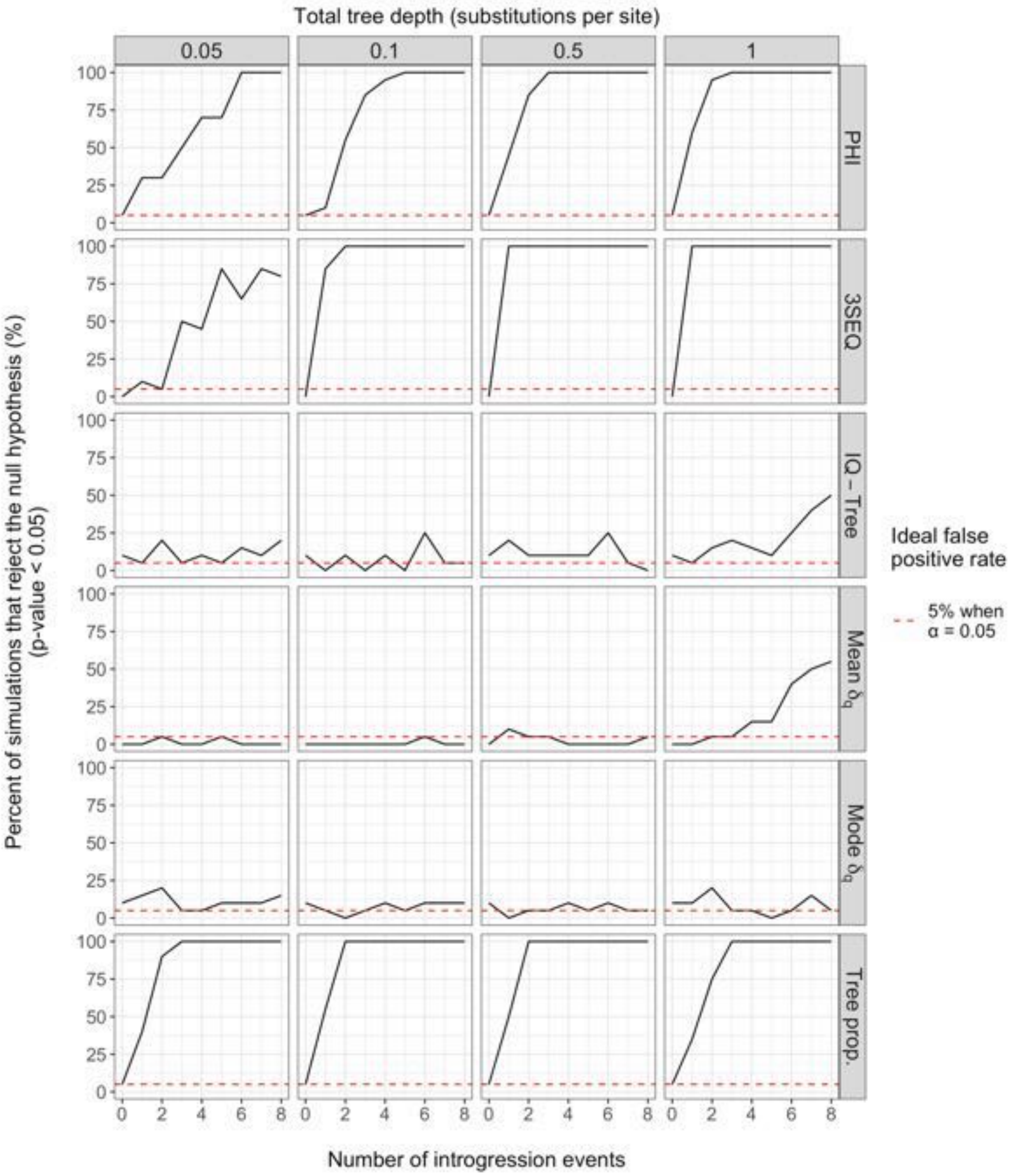


**Supplementary Figure 8:** p-values for an increasing number of nonreciprocal introgression events on a balanced 32-taxon tree with a tree depth of 0.5 substitutions per site. Each line represents the number of alignments (out of 100 replicates) that reject the null hypothesis of treelikeness. Each column shows a tree depth from 0.05 to 1 in substitutions per site. Each row shows results from a different test statistic. PHI is the p-value from the PHI test (implemented using PhiPack), 3SEQ is the p-value from the 3SEQ test, IQ-Tree is the p-value calculated using a parametric bootstrap from the proportion of resolved quartets (Likelihood Mapping), Mean $\delta_q$ and Mode $\delta_q$ are the p-values from the $\delta$ plotting method (calculated using a parametric bootstrap), and Tree prop. is the p-value for the tree proportion (calculated using a parametric bootstrap).
