## Supplementary Files README for "A test statistic to quantify treelikeness in phylogenetics"

A test statistic to quantify treelikeness in phylogenetics
Cherryh et al
- Supplementary Files README -

This folder contains supplementary figures and simulation data used for the analyses presented in the paper. Scripts to run analyses can be downloaded from https://github.com/caitlinch/treelikeness

# Supplementary figures
Contains supplementary figures showing results for event type (reciprocal and non-reciprocal) and tree age (0.1 - 1 substitution per site)

# Data files
For each of the two main analyses (increasing proportion of introgressed DNA and increasing number of introgression events) there are two files:
	* A test statistic file, containing the simulation parameters and test statistic results for the six tests
	* A p-value file, containing the simulation parameters and p-value results for the six tests
The column names include a reference to the six tests as follows:
	* "PHI" = Pairwise Homoplasy Index
	* "X3SEQ" = 3seq
	* "LM" = likelihood mapping
	* "delta_q" = delta plots
	* "tree_proportion" = tree proportion
	
